# A methodological framework for accommodating Cancer Genomics Information in OMOP-CDM using Variation Representation Specification (VRS)

**DOI:** 10.64898/2026.02.09.702490

**Authors:** Elisa Benetti, Guido Scicolone, Muhammad Tajwar, Corrado Masciullo, Gabriele Bucci, Michela Riba

## Abstract

The OMOP Common Data Model (OMOP CDM) in which observational health data are organized and stored is a broadly accepted data standard which helps clinical research facilitating federation study protocols. In case of cancer studies, there is a growing need to incorporate cancer genomics data in a standardized way.

Starting from a brief overview of the basic features of the OMOP CDM, we imagine a path of increasing complexity for including known biomarker genomic data coming from pathology or reports or clinical laboratory findings, towards storing thousands of known and unknown variants coming from genome sequencing data. Data should be stored using standardized identifiers, including those defined by the Global Alliance for Genomics and Health (GA4GH). We propose a scalable strategy for storing genomics variants in increasingly complex scenarios and present KOIOS-VRS, a pipeline that automates the conversion of VCF files into OMOP compatible format.

## INTRODUCTION

### OMOP Common Data Model

The Observational Medical Outcomes Partnership Common Data Model -OMOP CDM- (Observational Health Data Sciences and Informatics, 2021; Reinecke et al., 2021), a data standard developed by the Observational Health Data Sciences and Informatics (OHDSI), is a relational database consisting of standardized tables such as PERSON, MEASUREMENT, and OBSERVATION. [FIGURE 1] (Reich et al. (2024)

**FIGURE 1.**
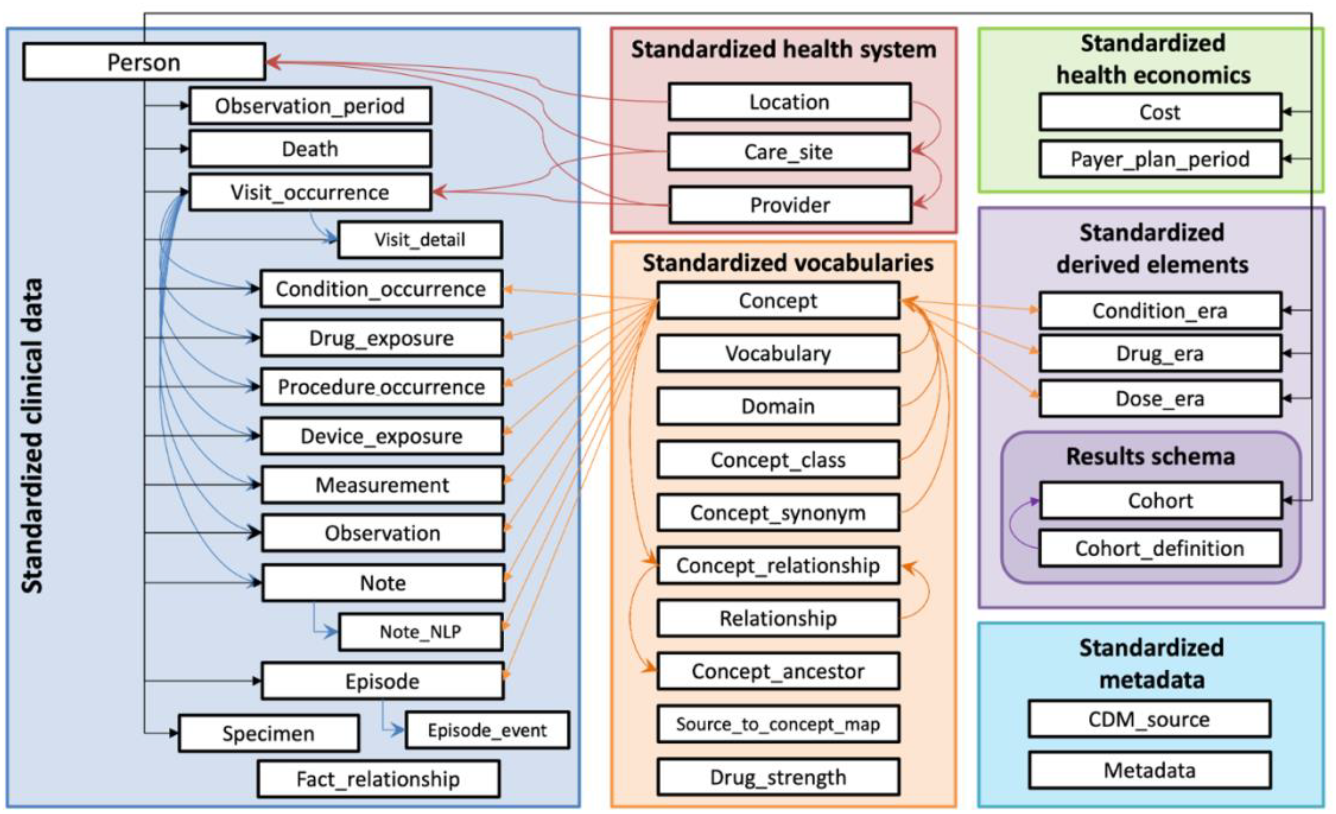
OMOP Tables

Conformance to the model requires that mandatory columns be filled using standard vocabularies (e.g., SNOMED, LOINC) accessible through resources like ATHENA. While the OHDSI Oncology Working Group has extended the model to support cancer conditions and treatments (Belenkaya et al., 2021), the representation of granular genomic variants remains an area of active development. Currently, Real World Data (RWD) genomic information is often stored as unstructured text, which significantly limits its utility for automated clinical research. In this work, we explore methods to bridge this gap by integrating GA4GH standards (Rehm et al., 2021) into the existing OMOP CDM framework or extension.

Each OMOP CDM table contains mandatory columns; among these the “*concept*.*id*^1^“ is of particular importance and should be filled using standardized OMOP vocabularies (Reich et al., 2024) [TABLE 1].

**TABLE 1.**
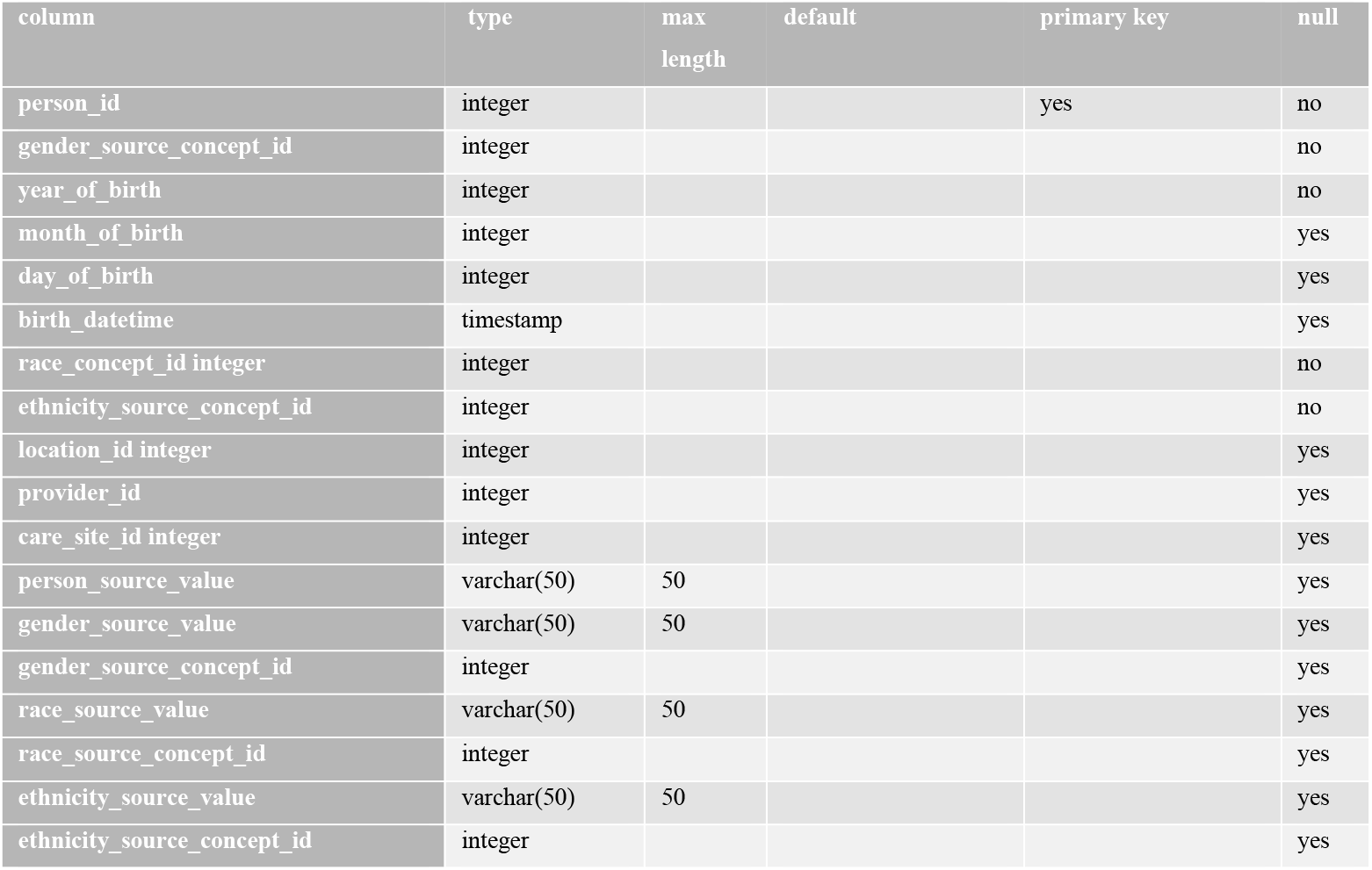
OMOP PERSON table with conformance specification^2^.

The ATHENA resource can be used to search and retrieve OMOP vocabularies [FIGURE 2] https://athena.ohdsi.org/search-terms/start

**FIGURE 2.**
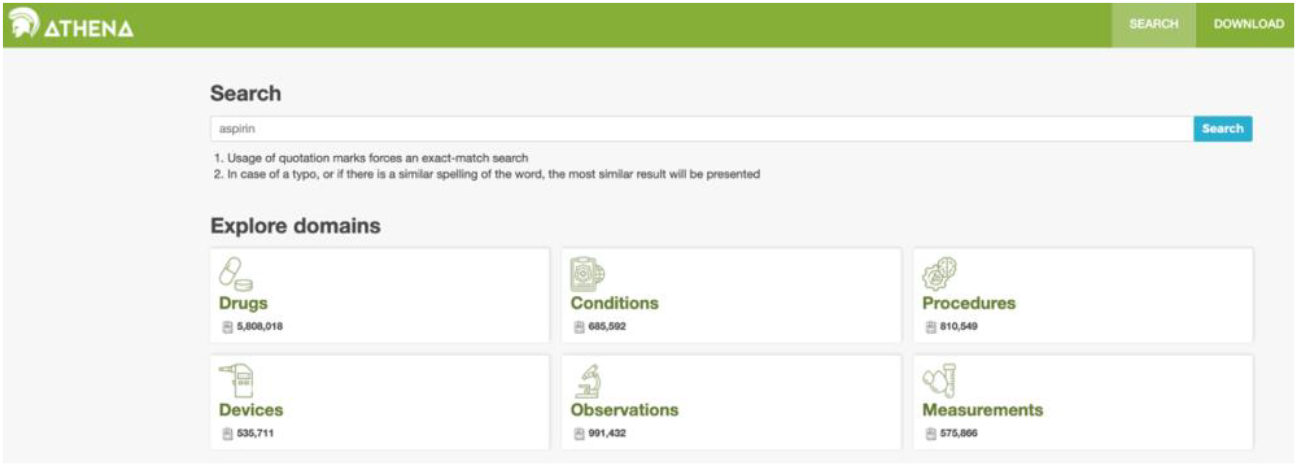
Athena.

### Genomic Information in OMOP

In the following sections, we outline several scenarios of increasing complexity for representing genomic data within the OMOP CDM.

#### Scenario1: USE STANDARD OMOP CDM TABLEs: MEASUREMENT FEW KNOWN BIOMARKERS MUTATIONS

As a first step, genomic information can be included as a biomarker (prognostic, diagnostic, therapeutic) in cancer patients’ data. This approach is suitable when only a limited number of well-characterized genomic markers are available.

In this scenario, genomic biomarkers can be included using the MEASUREMENT table of the OMOP CDM (https://ohdsi.github.io/CommonDataModel/cdm54.html#measurement) Each biomarker is encoded by assigning an appropriate LOINC code (https://loinc.org/) to the *measurement*.*concept*.*id*, integrated as OMOP *concept*.*id* [TABLE 2].

**TABLE 2.** BRAF code example. BRAF V600 biomarker is mapped to LOINC code to be inserted into the MEASUREMENT table.

| ID | CODE | NAME | CLASS | Concept | Validity | DOMAIN | Vocab |
| --- | --- | --- | --- | --- | --- | --- | --- |
| 616707 | 97025-1 | BRAF gene V600 mutations<br>[Identifier] in Tissue by Molecular<br>genetics method Nominal | Lab Test | Standard | Valid | Measurement | LOINC |

LOINC is a database providing standard codes in the domain of laboratory analytics with some expansions in pathology and genomics (Forrey et al., 1996).

The GA4GH Variation Representation Specification (VRS) is a computational framework designed for the deterministic and federated identification of genomic variants. It introduces a crucial paradigm by providing permanent and unique identifiers (e.g., ga4gh:VA.mJbj…) that allow variants to be identified consistently across different clinical and genomic datasets without relying on local database IDs.

We propose the integration of GA4GH VRS notation to encode the exact genomic variant associated with each biomarker. As illustrated in our framework, these identifiers can be successfully accommodated within the *value_source_value* column of the OMOP MEASUREMENT table. This approach is technically compatible with the current data model, as the standard VRS string (approximately 43 characters) fits within the 50-character limit of the target column [TABLE 3].

**TABLE 3.** OMOP Measurement Table.

| column | type | max length | default | primary key | null |
| --- | --- | --- | --- | --- | --- |
| measurement_id | integer |  |  | yes | no |
| person_id | integer |  |  |  | no |
| measurement_concept_id | integer |  |  |  | no |
| measurement_date | date |  |  |  | no |
| measurement_datetime | timestamp |  |  |  | yes |
| measurement_time | varchar(10) | 10 |  |  | yes |
| measurement_type_concept_id | integer |  |  |  | no |
| operator_concept_id | integer |  |  |  | yes |
| value_as_number | numeric |  |  |  | yes |
| value_as_concept_id | integer |  |  |  | yes |
| unit_concept_id | integer |  |  |  | yes |
| range_low | numeric |  |  |  | yes |
| range_high | numeric |  |  |  | yes |
| provider_id | integer |  |  |  | yes |
| visit_occurrence_id | integer |  |  |  | yes |
| visit_detail_id | integer |  |  |  | yes |
| measurement_source_value | varchar(50) | 50 |  |  | yes |
| measurement_source_concept_id | integer |  |  |  | yes |
| unit_source_value | varchar(50) | 50 |  |  | yes |
| unit_source_concept_id | integer |  |  |  | yes |
| value_source_value | varchar(50) | 50 |  |  | yes |
| measurement_event_id | bigint |  |  |  | yes |
| meas_event_field_concept_id | integer |  |  |  | yes |

We propose that the VRS notation can be accommodated into the *value*.*source*.*value* column of the MEASUREMENT table. This field refers to the original value of a variable as present in the clinical record (Electronic Health Record-EHR-for example). Example of a GA4GH VRS-notation:

`ga4gh:VA.mJbjSsW541oOsOtBoX36Mppr6hMjbjFr`

`“sequence_id”: “ga4gh:SQ.hSeY-byldamzZ5GGrmsCvAoco-nvCXYe”`

https://search.cancervariants.org/api/v1/associations?size=10&from=1&q=ga4gh:VA.yYu4TSKSJN9MUdJHVbiVctgvev0SkuOM

We evaluated the feasibility of storing such VRS identifiers within individual OMOP CDM table^3^ and confirmed that this approach is technically compatible with the data model.

| column | type | Max length |
| --- | --- | --- |
| value_source_value | varchar(50) | 50 |

#### Scenario2: Larger panels of mutations recorded in OMOP^4^ A NUMBER OF KNOWN MUTATIONs

This scenario addresses the representation of larger panels of known genomic mutations, which may not necessarily be reported in pathology reports for a specific disease but are nonetheless clinically relevant.

From a technical point of view to specify an OMOP measurement we need to include a concept.id which is mandatory, we imagine using the OMOP genomic vocabulary which contains now >100,000 entries^5^ and accommodate the VRS code inside the *value*.*source*.*value* column of the MEASUREMENT table.

#### Scenario3: large number of mutations, including Variants of Unknown Significance (VUS)

In the previous scenarios we imagined a way to store small or medium number of known mutations, annotated in OMOP with LOINC or OMOP genomics vocabularies. We now imagine how we could store potentially unknown mutations in a way suitable for OMOP rules. We need an identifier to be able to constitute the OMOP entry: *concept*.*id*

We propose to use a generic one defining a Variant of Uncertain Significance (VUS) by means of a LOINC code^6^.

We propose to use a generic one defining a VUS https://athena.ohdsi.org/search-terms/terms/1028197

We can then accommodate all the same the VRS code in the *value*.*source*.*value* column^7^

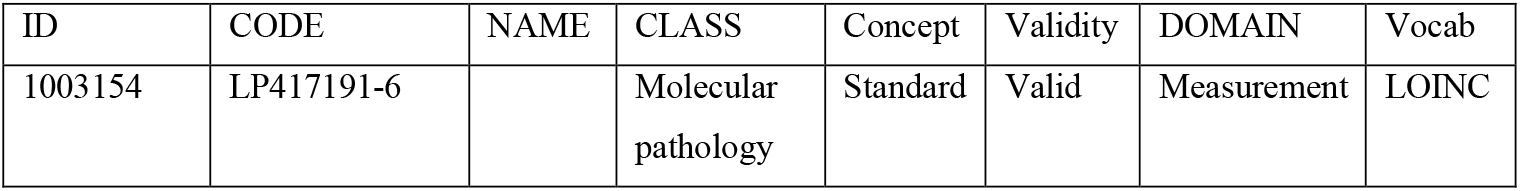

## MATERIALS AND METHODS

To define the strategy to include genomic data into OMOP we reviewed the current CDM v5.4 (https://ohdsi.github.io/CommonDataModel/cdm54.html) with specific focus on the MEASUREMENT TABLE specifications and requirements due to the operative derivation of data from molecular laboratory sources, such as for example biochemical tests and results, usually stored in this table. We then examined the available OMOP vocabularies to describe genomics data with deeper insight into the OMOP GENOMIC, available for download at Athena search tool (https://athena.ohdsi.org/vocabulary/list).

We evaluated and used the KOIOS tool that, starting from variants in a VCF file, can provide a mapping to OMOP GENOMIC vocabulary (https://github.com/OHDSI/Koios).

To additionally support the unique variant representation defined by the GA4GH VRS standard, we reviewed the VRS specifications (https://www.ga4gh.org/product/variation-representation/) and developed the R KOIOS_VRS package, that extends the KOIOS workflow (https://github.com/gbucci/koios_vrs). Starting from a VCF file, KOIOS_VRS produces both the corresponding OMOP genomic record, which can be included in the OMOP MEASUREMENT table, and the VRS-compliant variant identifiers via the GA4GH registry API (https://reg.genome.network/).

The package outputs the VRS in GA4GH Phenopackets standard (https://phenopacket-schema.readthedocs.io/en/latest/variant.html) in the “*variation*” field.

To face the problem of scaling up to include hundreds or thousands of variants for a single patient and to store genomic information in a standard format we studied the proposed extension OMOP G-CDM (https://github.com/OHDSI/Genomic-CDM) and worked on a simplified table.

Tests have been conducted on VCFs derived from synthetic data (https://zenodo.org/records/15754326).

## RESULTS

### KOIOS-VRS pipeline

To facilitate the transition from raw data to a standardized CDM, we developed KOIOS-VRS, a multi-platform genomic annotation pipeline which works in the following steps [FIGURE 3].

**FIGURE 3.**
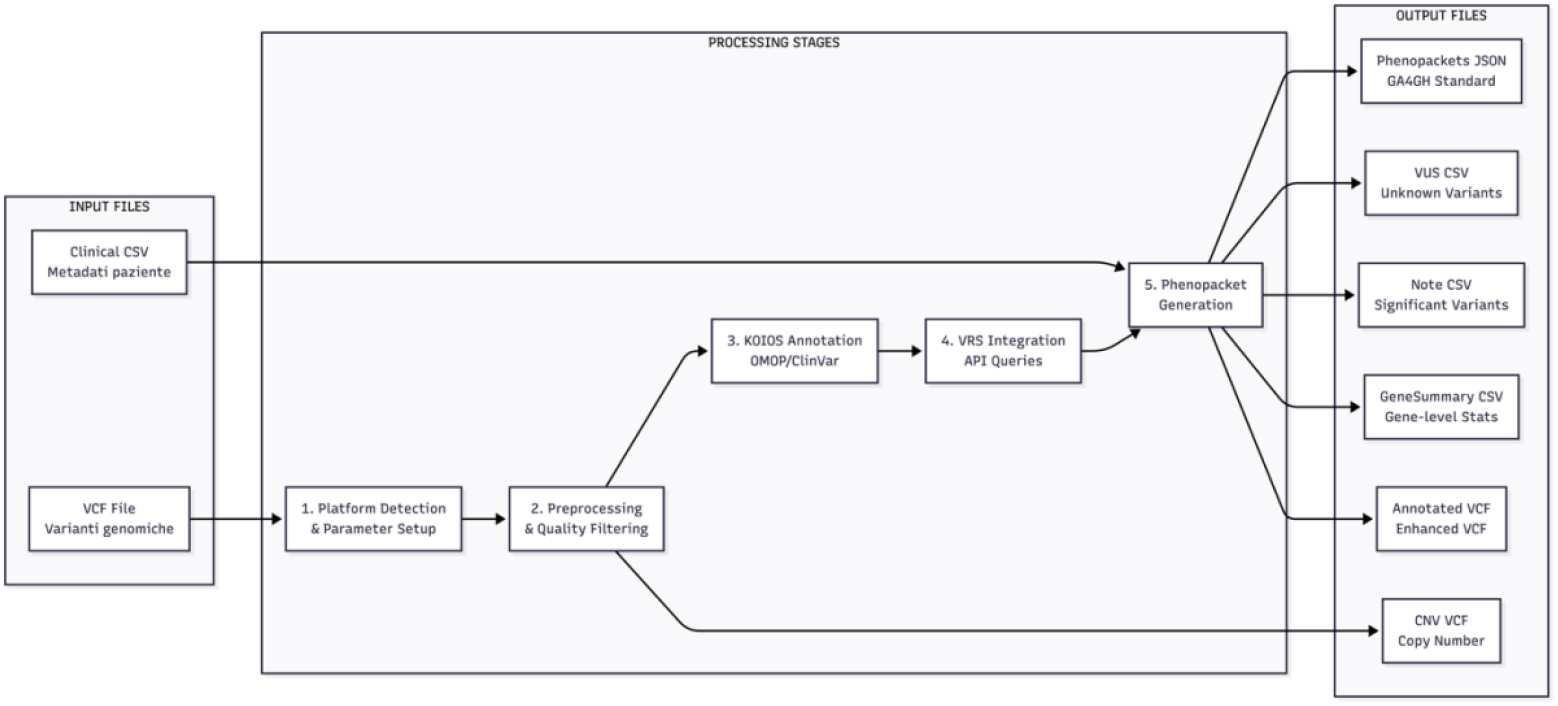
KOIOS-VRS flowchart.

#### 1. Data Input and Platform Detection

The pipeline accepts VCF files (v4.1+) and clinical metadata in CSV format. A key feature is the automatic detection of sequencing platforms (e.g., Ion Torrent, Illumina) from the VCF header. This allows the tool to apply optimized quality filters, such as Allele Frequency (AF) and Depth of Coverage (DP) thresholds specific to each technology.

#### 2. Standardization and VRS Integration

The pipeline leverages the KOIOS logic to automate the conversion of Raw Variant Call Format (VCF) into standardized Human Genome Variation Society (HGVS) notations. This step is critical as it provides the necessary nomenclature to map variants directly to the OMOP genomic vocabulary concept identifiers, ensuring consistency across clinical and genomics datasets (OHDSI. Koios: OMOP-based genomic variant annotation. Available from: https://github.com/OHDSI/Koios).

The tool KOIOS has been developed to produce, starting from a Variant Call Format (VCF) file the corresponding Human Genome Variation Society (HGVS)^8^ codes, which can be matched with OMOP genomic vocabulary concept_id, HGVS code can be used to produce a corresponding VRS (Wagner et al., 2021) code by means of matching with pre-existent collections^9^ or producing the code from the HGVS notation by means of a software tool^10^. To ensure permanent and unique identification, the pipeline integrates VRS (Variant Registration Service) identifiers. These identifiers can be accommodated within the *value_source_value* column of the OMOP MEASUREMENT table.

The KOIOS-VRS pipeline generates several structured outputs to support both clinical and bioinformatic needs:

- **Phenopackets JSON**: Phenopacket standard representation (Jacobsen et al., 2022), which includes both patient clinical data and variant interpretations with VRS IDs and OMOP mappings.
- **Annotated VCF**: Incorporates VRS identifiers and OMOP concept IDs directly into the INFO field.
- **Clinical and VUS CSVs**: Separate files for clinically significant variants (“Known Variants”) and VUS, facilitating refined clinical interpretation.

Our preliminary tests illustrate that the pipeline can successfully process hg19 and hg38 data, performing automatic liftover when necessary. The use of VRS identifiers provides a robust way to store complex genomic data within the 50-character limit of standard OMOP columns like *value_source_value*.

#### Use case: Synthetic integration of cancer genomic variants into OMOP-CDM using VRS Use case description

To demonstrate the practical applicability of the proposed methodological framework, we present a fully synthetic use case illustrating how cancer genomic variant information can be represented using the GA4GH Variation Representation Specification (VRS) and accommodated within the OMOP Common Data Model (CDM). This example is provided solely for methodological illustration and does not involve real-world patient data.

#### Simulated Real World Scenario

Hereafter we present a simulated example. All identifiers, clinical events, and genomic attributes used in this scenario are synthetic and were generated exclusively for illustrative purposes.

For a hypothetical oncological female subject, diagnosed with melanoma at the age of 60 years, the sequencing reported several somatic mutations. A somatic *BRAF* p.Val600Glu (V600E) variant, a well-characterized alteration frequently reported in cancer genomics, was identified. The raw VCF file, obtained from sequencing and standard variant call pipeline, and the associated clinical data in tabular format, can be used as input to the KOIOS-VRS pipeline (Supplementary information: melanoma_sample.vcf and melanoma_sample.csv). The output of the pipeline on the synthetic example above gives the following results (Supplementary information: melanoma_final_Phenopackets.json).

#### VRS-based representation

The genomic variant is represented as a “VRS Allele” object defined by a precise sequence location on chromosome 7 (GRCh38) and a single nucleotide substitution corresponding to the V600E amino acid change. A synthetic VRS allele identifier is computed to enable unambiguous, computable reference to the variant across systems since the VRS representation provides a normalized, schema-independent description of the variant, decoupled from local database implementations.

#### Accommodation within OMOP-CDM

Following the proposed framework, the VRS allele is accommodated within the OMOP-CDM using existing standardized tables without modification of the core data model. Specifically:

- The genomic variant is recorded as a “measurement” entry in the OMOP MEASUREMENT table.
- The VRS allele identifier is stored in the measurement_source_value field to preserve precise genomic identity.
- Standard OMOP vocabulary concepts are used to annotate the variant, while VRS remains the primary computable identifier.
- Additional contextual attributes, including reference genome assembly and variant type, are preserved through structured metadata as defined by the framework.

#### Illustrative OMOP-CDM record

Hereafter a simplified synthetic OMOP-CDM representation of the genomic variant (TABLE 4).

| person_id | measurement_concept_id | measurement_source_value | measurement_date | value_as_concept_id |
| --- | --- | --- | --- | --- |
| 100001 | 36010025 | {<br>"id": "variant_interpretation_1",<br>"variantInterpretation": {<br>"id": "SAMPLE_ID_M1_7_140753336",<br>"vrsAllele": {<br>"id": "ga4gh:VA.HaPTmn-<br>rrjRoZnIVw1l4AZPa6YHa2ojh",<br>"type": "Allele"}<br>}<br>} | 2025-01-01 | BRAF:V600E |

All values shown are synthetic and do not correspond to real individuals or clinical records.

#### Accommodation within Phenopackets standard

Clinical and genomic variation data are represented in a Phenopackets standard file (supplementary_information.pdf, melanoma_final_Phenpackets.json)

#### Demonstrated utility

This use case illustrates how the proposed framework:

1. Enables precise and interoperable representation of cancer genomic variants using VRS;
2. Integrates genomic information into OMOP-CDM without introducing custom tables or non-standard extensions;
3. Supports reuse of genomic data across OMOP-based analytical workflows while maintaining semantic clarity.

## DISCUSSION AND PERSPECTIVES

In the previous sections we have shown approaches for representing genomic variants within the OMOP CDM standard tables facing the issues of accommodating an increasing number of mutations, both known and unknown. We demonstrate the possibility to integrate in OMOP, a widely used standard clinical database format, the VRS, the best suited codification of genomic variation, as defined by the GA4GH Consortium.

We spotted a potential dimensionality issue related to the nature of genomic data. Consequently, we explore the possibility to include data of higher orders of magnitude, for example the information contained in VCF coming from whole exome sequencing (WES) or even whole genome sequencing (WGS), which contain thousands to millions of variants per patient. This volume of data poses a significant scalability challenge as it differs from typical OMOP measurement data, which are designed to represent discrete clinical observations or laboratory results.

### Extensions to the OMOP CDM

#### OMOP Genomic Common Data Model (G-CDM)

A genomic extension for the OMOP CDM, known as the OMOP Genomic Common Data Model, G-CDM, has been proposed to support the structured representation of high-throughput genomic data.

The G-CDM includes dedicated tables, like VARIANT_OCCURRENCE and GENOMIC_TEST, for the storage of genomic information along with the keys needed to bridge with the core OMOP tables [FIGURE 4]. The G-CDM offers an extensive schema to detail genomic variation proposing a lot of mandatory and non-mandatory variables, but we observed that its widespread adoption within the OHDSI community is still evolving.

**FIGURE 4.**
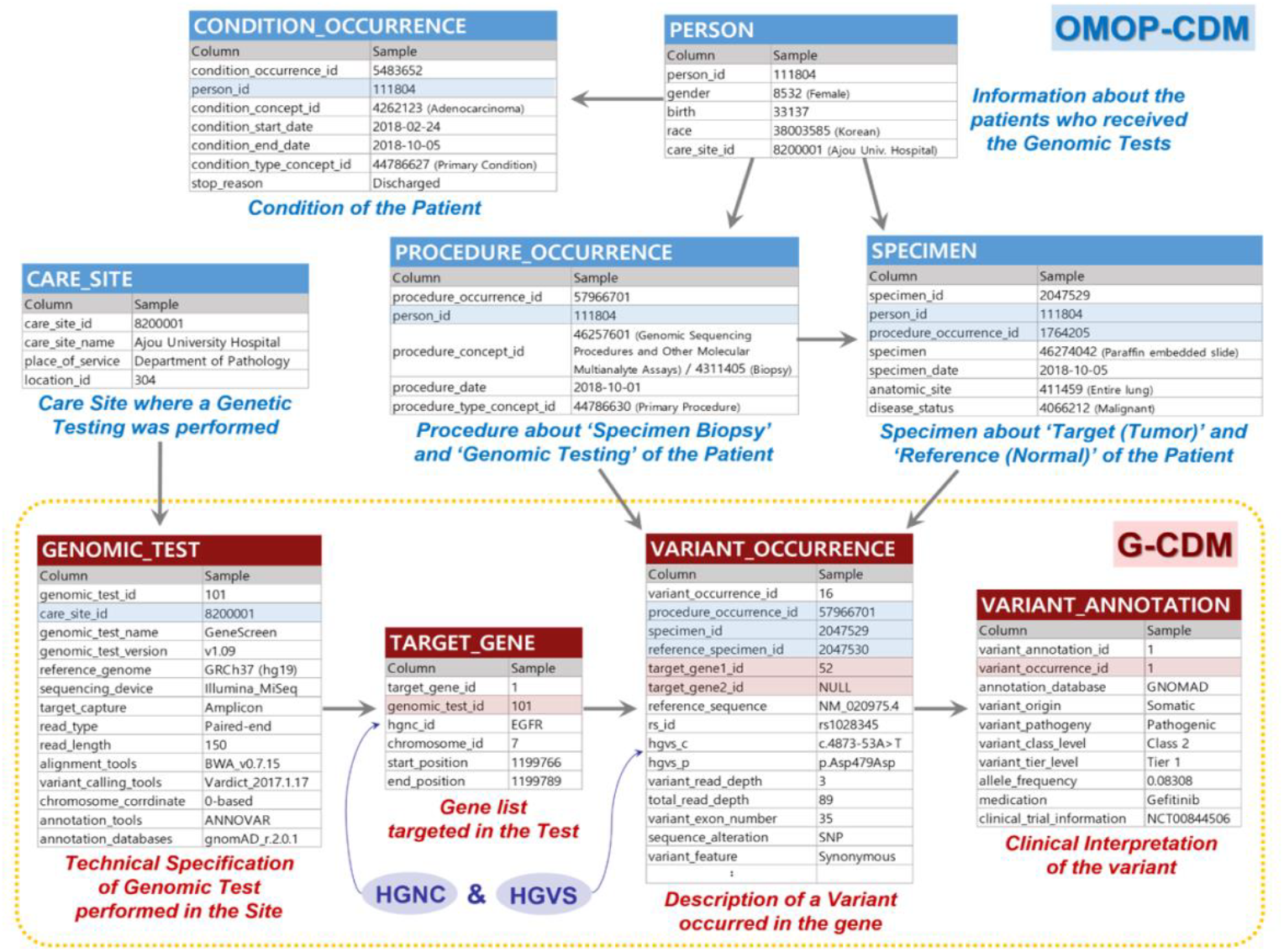
G-CDM, genomic CDM.

The full G-CDM specification and implementation details are publicly available at: https://github.com/OHDSI/Genomic-CDM

#### A Scalable Proposal: The “GENE” table

We propose an alternative and simplified approach, if compared to the G-CDM, for the OMOP model by including a single “GENE” additional table dedicated to genomic results. In this approach, selected column formats could be defined with additional variable types, for example JSON format, to better store GA4GH VRS codes and related metadata. This design has the potential to:

-enhance scalability by grouping variant data into a flexible JSON structure rather than multiple relational rows.

- improve the analytical workflows after database creation, by leveraging JSON-specific database libraries to filter and retrieve specific variants across large cohorts.

This proposal represents an initial conceptual framework and that we hope will contribute to the ongoing discussion within the OHDSI community regarding the optimal balance between relational structure and genomic data volume.

## CONCLUSIONS

Integrating GA4GH standards like VRS into the OMOP CDM is a promising step toward interoperable cancer research. The KOIOS-VRS pipeline describes a functional bridge between raw genomic data and standardized clinical databases. Future efforts should focus on refining these storage strategies to support the full scale of precision oncology data within the OHDSI community.

## Data Availability

The KOIOS-VRS tool is open-source and available at: https://github.com/gbucci/koios_vrs

## Supplementary Information

The Supplementary information file contains input and output test data including VCF, clinical metadata (csv), and json file. (supplementary_information.pdf, melanoma_sample.vcf, melanoma_sample.csv and melanoma_final_Phenopackets.json)

## Ethics Statement

This work is based on methodological considerations and illustrative examples. No real-world patient data were used, and ethical approval was not required.

## Acknowledgments

This work was supported by the DigiOne I3 WP4 (Digital Oncology Innovation Initiative https://www.digione-i3.eu/) and the IRCCS Ospedale San Raffaele.

We would like to thank for fruitful discussion Arcangela De Nicolo, Aurora Maurizio, Giorgia Gandolfi, Francesca Genova, Cinzia F. Sala, Marco J. Morelli and Giovanni Tonon

## Footnotes

1 In FIGURE 2 as an example gender_concept_id or ethnicity_concept_id

2 Conformance defines if a variable is mandatory or not, in this example the mandatory ones cannot assume the NULL value

3 The VRS json code can be accomodated into the *value*.*source*.*value* column:

4 Some mutations may be not even been reported in the pathology reports

5 In this second scenario we imagine to scale up from a small number of biomarker mutations (less than 10) to the content cancer focused gene set, let’s say less than 100 mutations

6 There exist other options to classify in OMOP a VUS, e.g. the following, anyhow it is better to use standard and valid, the following is a non standard code https://athena.ohdsi.org/search-terms/terms/1028197

7 The VRS code has to be calculated from the HGVS code of the mutation

8 HGVS code is used to describe variants in DNA, RNA, and protein sequences. It is a recognized international standard for communicating gene variations in clinical reports and sharing them in publications and databases.

9 Some database storing already documented mutations in VRS fomat and HGVS codes are available and under evaluation

10 A software tool do exist for producing the VRS notation from a HGVS, https://github.com/ga4gh/vrs-python

## Notes

https://github.com/gbucci/koios_vrs

### Competing Interest Statement

The authors have declared no competing interest.

