## Supplementary information for "A methodological framework for accommodating Cancer Genomics Information in OMOP-CDM using Variation Representation Specification (VRS)"

### *Simulated Real World Scenario*

Input VCF File: melanoma_sample.vcf, available at:

https://github.com/gbucci/koios_vrs/blob/main/examples/melanoma_sample.vcf

Input Clinical Metadata CSV: melanoma_sample.csv, available at:

https://github.com/gbucci/koios_vrs/blob/main/examples/melanoma_sample.csv

Output Phenopackets JSON: melanoma_final_Phenopackets.json, available at

https://github.com/gbucci/koios_vrs/blob/main/melanoma_final_Phenopackets.json
